# Deep learning models reading clinical data and liver omics strongly distinguish NASH from steatosis and suggest new genes involved in liver disease severity

**DOI:** 10.1101/2025.10.10.681581

**Authors:** Nicolas Gambardella, Smaïn Fettem, Mathilde Boissel, Lijiao Ning, Violeta Raverdy, Marwa Afnouch, Souhila Amanzougarene, Mehdi Derhourhi, Bénédicte Toussaint, Emmanuel Vaillant, Amna Khamis, Philippe Lefebvre, Bart Staels, François Pattou, Philippe Froguel, Amélie Bonnefond

## Abstract

**Background & Aims:** Metabolic dysfunction-associated steatotic liver disease (MASLD, previously NAFLD) is a frequent co-morbidity of obesity and diabetes, with prevalence increasing worldwide in all age groups and both sexes. Only early stages of the disease are fully reversible. Recognising liver disease stages and elucidating the molecular underpinning of their progression are thus medically important. We developed a deep learning model to recognise simple steatosis from steatohepatitis combining liver transcriptomics, epigenetics, and clinical data.

**Methods:** We used clinical data, liver gene expression and liver DNA methylation gathered from 300 patients with obesity of the ABOS cohort (80 without NAFLD, 137 with simple steatosis, 83 with steatohepatitis). We selected non-redundant clinical variables, gene expressions and CpGs methylation levels most associated with severity using unsupervised approaches. We designed a multi-module, multi-layer perceptron to predict patients’ liver status. We trained five model instances on independent training/test sets and combined the predictions.

**Results:** We used a score based on gene expression/DNA methylation and relevant principal component analysis (PCA) loadings to select 200 genes and 260 CpG methylations. Models trained on the three modalities reached an AUC of 0.945 overall on a validation set with accuracies above 81% for simple steatosis and 88% for NASH, outperforming any other machine learning model so far. We retrieved patient clusters previously found using clinical variables in the latent space of our clinical data module, but not in the gene expression and DNA methylation modules. While all three modules are needed to reach the best prediction accuracy in all classes, the gene expression module had the most impact on the decision. Independent models weighted gene expression inputs similarly, shining light on their importance. The most impactful genes were linked to immune responses and extracellular matrix. However, many of those genes were previously unassociated with steatotic liver disease onset or progression.

**Conclusions:** A multi-omics deep-learning model can recognise steatohepatitis from simple liver steatosis with an AUC of 0.945 and identify new genes potentially involved in NAFLD progression. Gene expressions profiles predicting disease severity are largely different from those specific of clinical variable clusters.

**Impact and implications:** This study suggests that clinical variables are not sufficient to recognise the severity of steatotic liver disease with high accuracy, but model efficiency increases when used together with liver epigenetics and transcriptomics.

## Introduction

Metabolic Dysfunction-Associated Steatotic Liver Disease (MASLD), formerly Non-Alcoholic Fatty Liver Disease (NAFLD), encompasses a range of liver conditions characterised by excessive fat accumulation in the liver (Targher *et al*, 2025). These conditions are most often co-morbidities of obesity, type 2 diabetes, and dyslipidemia. MASLD initiates as relatively benign and reversible steatosis (MAFL, formerly NAFL, also called fatty liver) but may progress into severe forms of liver diseases, including steatohepatitis (MASH, formerly NASH, presenting inflammation), fibrosis, and cirrhosis, and hepatocellular carcinoma (Lekakis & Papatheodoridis, 2024). While the official nomenclature is now MASLD and MASH, there are differences in diagnosis between NAFLD and MAFLD (Eslam *et al*, 2020) and then MASLD (Rinella *et al*, 2023). In this paper, we will keep the terminology NAFLD, NAFL, and NASH in the analyses since those were the diagnoses made when the data was collected.

The burden of MASLD is increasing rapidly, already affecting 30% of adults, largely driven by the global increase in the prevalence of metabolic diseases (Miao *et al*, 2024). While isolated liver steatosis and even moderate MASLD are reversible upon weight loss consecutive to diet, bariatric surgery (Fakhry *et al*, 2019; Lassailly *et al*, 2015) or treatments, such as resmetirom (Harrison *et al*, 2024), or GLP-1 and other incretin receptor activators (Sanyal *et al*, 2025; Loomba *et al*, 2024; Sanyal *et al*, 2024), this is not the case once bridged fibrosis and cirrhosis appear. Therefore, it is crucial to recognise when these irreversible liver diseases are present, and to understand the molecular mechanisms underpinning the evolution from steatosis to steatohepatitis, as this may significantly impact patient’s prognosis and care. Recently, clusters of patients with different characteristics and possibly different disease trajectories have been proposed, mainly based on clinical data (Raverdy *et al*, 2024), but without multi-omic analysis.

Currently, accurately distinguishing between inflamed and non-inflamed steatotic livers requires hepatic biopsies, which pose significant risks for patients. In addition, the high degree of cellular heterogeneity of a sick liver means that such a histopathology diagnosis might be stochastic. Non-invasive biomarkers and panels of biomarkers have thus been developed over the years, presenting variable accuracies and limitations (Piazzolla & Mangia, 2020; Sanyal *et al*, 2023; Zoncapè *et al*, 2024). The combination of statistical measures to obtain a score is generally quite simple, such as a weighted sum fed to a logistic regression. In addition, modern architectures are now appearing to make the most of non-invasive procedures such as ultrasounds (Guo *et al*, 2024). However, although such approaches can measure the extent of liver disease, they cannot distinguish between simple steatosis and steatohepatitis. Deep learning offers means to integrate diverse data types and use complex non-linear and non-monotonic relationships. Various deep learning models have been proposed to diagnose and score MASLD (Jimenez Ramos *et al*, 2024). As for other diseases, convolutional neural networks have been used for a few years in histopathology to recognise features on biopsy slides (Heinemann *et al*, 2019, 2022). Using deep learning with omics datasets, such as gene expression profiles, is more recent (Park *et al*, 2023; Oh *et al*, 2024).

We developed a deep learning model that combines clinical measurement, liver gene expression and DNA methylation to distinguish between NAFL and NASH in 300 patients with severe obesity. The analysis of the features prioritized by the model provides leads toward new molecular networks underpinning the disease progression.

## Materials and methods

### Dataset

The Atlas Biologique de l’Obésité Sévère (ABOS) cohort is a prospective multi-tissue cohort based in Lille (France), with samples collected from individuals during bariatric surgery (clinicaltrials.gov, NCT01129297). All patients provided written informed consent before inclusion. The Comité de Protection des Personnes Nord Ouest VI (Lille, France) granted the study’s ethical approval. The present study focused on the data from 300 ABOS participants of European ancestry. Sixty-six demographic characteristics, anthropometric measurements, medical history, medication use, and clinical laboratory test data were prospectively collected before or at the time of surgery. Histopathological scales were used to characterise Steatosis (quantitative, [0-3]), Ballooning ([0-2] = [none, some, much]), and Inflammation (number of foci, [0-3]). 80 patients were characterised as healthy (S = 0, B = 0, I = 0) and 83 as suffering from steatohepatitis (S > 0, B > 0, I > 0). The remaining 137 patients were classified as suffering from steatosis but no steatohepatitis. Throughout the paper we label patients without liver steatosis as “healthy”, which by no means implies that those individuals did not display any unrelated disorders. They were just classified as healthy by opposition to simple steatosis and steatohepatitis.

### Clinical variables

Some of the 66 variables were alternative versions with different units, non-disjoint measurements (e.g., high-density lipoproteins (HDL), low-density lipoproteins (LDL), and total cholesterol), derived variables (e.g., averaged fasting glycaemia is the mean of glycaemia measured in isolation or at time 0 of an Oral Glucose Tolerance Test (OGTT), homeostasis model assessment of insulin resistance [HOMA2 IR] is derived from OGTT and fasting insulin). Finally, the current study focussed on the molecular underpinning of NAFLD, and we ignored the treatments. We retained 16 non-redundant laboratory analyses relevant for liver disease for which most patients had records, plus the sex and patient age. The dataset still exhibited a fair number of missing values. In order to keep a dataset as large as possible we imputed them. Two subjects, one healthy and one with steatosis, missed the same nine variables and were removed. 12% of the ultrasensitive C-reactive protein (CRP) values were missing. However, considering the importance of CRP as a marker for inflammation, we kept *CrPus* (ultrasensitive CRP) and imputed the missing values. We imputed missing values with the k nearest neighbours (kNN) method of the *VIM* R package v6.2.2 using k of 5 (Kowarik & Templ, 2016). Values were imputed for (see Table 1 for variable names) fasting insulinaemia (*fast_insul*, 15 values), glycated hemoglobin (*hba1c*, 2 values), LDL cholesterol (*LDL*, 2 values), α2 macroglobulin (*a2_macroglob*, 3 values), haptoglobin (*haptoglobin*, 2 values), Apolipoprotein A1 (*apoA1*, 1 values), Ultrasensitive CRP (*CrPus*, 32 values), lymphocytes (*lymphocytes*, 2 values). All the PCAs in the article were performed with the *prcomp* function of the *stats* R package v4.4.2, the variances being obtained with the *factoextra* package v1.0.7.

**Table 1.**
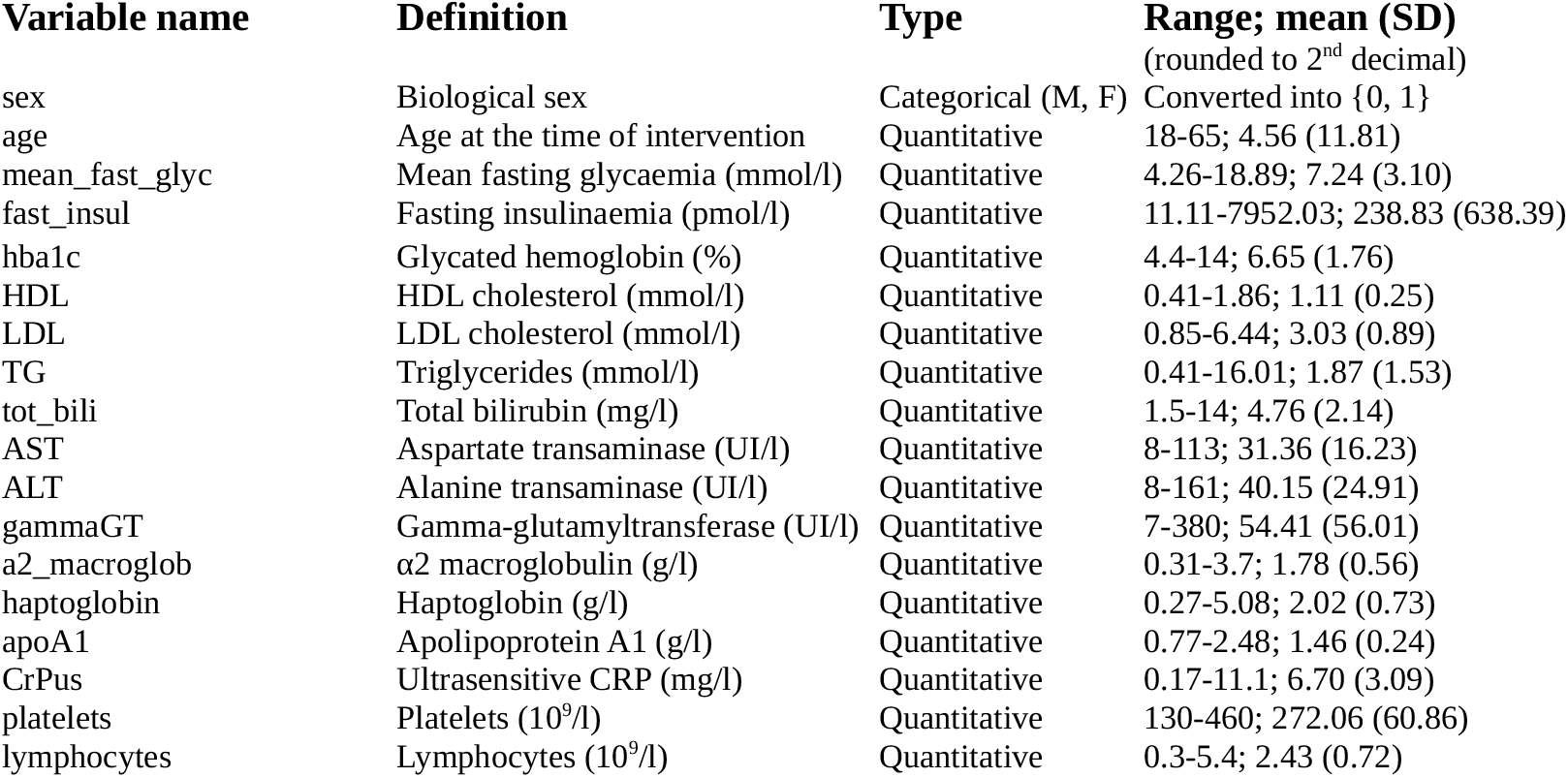
List of anthropometric and medical variables used in the models.

### Transcriptomics

RNA was extracted from liver biopsies, and total RNA-sequencing was performed on 200 ng of RNA using the KAPA RNA HyperPrep Kit with RiboErase HMR (Roche Sequencing). The libraries were sequenced in 2×75 bp paired-end reads using the NovaSeq6000 Illumina system.

Demultiplexing of sequence data was performed using Illumina’s bcl2fastq Conversion Software v2.20.0.422. The QC was performed using *FastQC* v0.11.9. The removal of adaptor sequences and low-quality bases was performed with *Trimmomatic* v0.39 (Bolger *et al*, 2014). Sequence reads were then mapped to the human genome (hg38) using *Star aligner* v2.7.3a (Dobin *et al*, 2013). On average, 72 M reads per sample were generated and 95 % were accurately mapped. Transcripts were quantified using *RSEM* v1.3.0 (Li & Dewey, 2011). Differential expression analysis was performed with *DESeq2* v1.42.1 and gene enrichment analysis with *WebGestalt 2024* (Elizarraras *et al*, 2024). The transcript expression score used for selecting genes to feed the multi-layer perceptrons (MLPs) was:

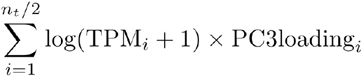

Where *i* is the number of genes selected at both extremities of the ordered PC3 loadings and *n*_*t*_ is the total number of transcripts. We established thresholds with the *glm* function, *category* based on the score, and the binomial family.

### DNA methylation

DNA was extracted from liver biopsies collected during bariatric surgery for all individuals using the DNeasy Blood & Tissue kit (Qiagen, Germany). Bisulfite conversion of 800 ng of genomic DNA was performed using the EZ-96 DNA Methylation kit (Zymo Research) following the manufacturer’s protocols, and bisulfite-converted DNA was subjected to a genome-wide DNA methylation analysis performed using Illumina’s Infinium 850K Methylation EPIC BeadChip array (Illumina, Inc., San Diego, CA, USA). All samples were randomised across the chips and analysed on the same machine by the same technician to reduce batch effects. The resulting DNA methylation IDAT files were imported using the *minfi* R package (Aryee *et al*, 2014) for further processing and quality control (QC). The following probes were excluded from further analysis: probes with a detection p-value greater than or equal to 0.01 for at least one sample, cross-hybridizing probes, probes with a bead count less than 3 in at least 5 % of the samples, non-CpG probes, and probes which lie near single nucleotide polymorphisms (SNPs), probes on chromosomes X and Y. The resulting dataset contained the methylation of 737 821 CpGs.

We computed M values (Du *et al*, 2010) as:

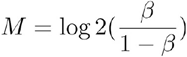

where β represent the measured methylation beta values. We obtained age-corrected M values by fitting linear regressions for each CpG in function of age, as:

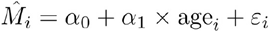

where *α*_*0*_ and *α*_*1*_ are parameters, and keeping the residuals *ε*_*i*_. The DNA methylation score used for selecting CpGs to feed the MLPs was:

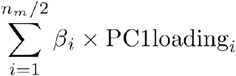

Where *i* is the number of CpGs selected at both extremities of the ordered PC1 loadings and *n*_*m*_ is the total number of methylation sites. We established thresholds with the *glm* function, *category* based on the score, and the binomial family.

### Multi-layer perceptron

We wrote the models in Python with *Tensorflow/Keras* v2.12. The clinical data module takes as input 16 laboratory measurements, age and sex. After dropout (20%), the data was fed to fully connected layers of 10 hyperbolic tangent neurons (tanh), a 50% dropout layer and 5 sigmoid neurons. Input data was fed raw since the normalisation was done within the models after the dropout layer. The RNA-seq module takes as input the transcript per million reads (TPM) expression of 200 genes. After dropout (20%) and normalising layers, data was fed to fully connected layers of 50 and 10 hyperbolic tangent neurons, each followed by a 50% dropout layer and 5 sigmoid neurons. The CpG methylation module takes as input the methylation beta-values of 260 CpGs. After dropout (20%) and normalising layers, data was fed to fully connected layers of 50 and 10 hyperbolic tangent neurons, each followed by a 50% dropout layer, and 5 sigmoid neurons. We concatenated the outputs of each module and fed it to a fully connected layer of 5 sigmoid neurons and a 50% dropout layer. The initial 20% dropout layers were followed by a whole set normalising layer, and each 50% dropout by a batch normalising layer. The output was provided by a layer of 3 softmax neurons. An L2 kernel regularizer was applied to each dense layer of the individual modules. We trained the models using a categorical cross-entropy loss function, an Adam optimiser (initial learning rate of 0.001), and an accuracy metrics. The training lasted 10 000 epochs, with an early stopping after a patience of 2000 epochs and a minimum delta accuracy of 0.001. We used shuffled batches of 16 subjects. We computed an overall prediction score for each class as the mean of the predictions from the five models. Receiver operating characteristic (ROC) curves were plotted with the R package *plotROC* v2.3.1 (Sachs, 2023) and areas under the curve (AUC) were computed with the package *pROC* v1.18.5 (Robin *et al*, 2011). Multi-class accuracy was the sum of true positives for the three classes divided by all predictions. Overall sensitivity, specificity, and precision were the means of those descriptors over the three classes. The clustering of latent coordinates was done with the *heatmap*.*2* function of the *made4* R package v1.78. UMAPs of patient embeddings in latent spaces were obtained with the native R algorithm of the *umap* R package v0.2.10, using 10 neighbours.

### Gene interaction inference

As in previous publications (Gambardella *et al*, 2019; Montalban *et al*, 2022, 2024), we inferred gene interactions by combining the results of *Context Likelihood of Relatedness* (CLR) (Faith *et al*, 2007), a mutual-information-based approach providing undirected edges provided by the R package *minet* v3.60 (Meyer *et al*, 2008), and *GEne Network Inference with Ensemble of trees* (GENIE3) v1.24 (Huynh-Thu *et al*, 2010), a tree-based regression approach providing directed edges. Both methods were the best performers in their category at the DREAM5 challenge (Marbach *et al*, 2012). We retained only the non-zero edges involving one of the 200 genes used as input to the RNA-seq MLP module (3162288 of the 584358193 edges) and we used the R package *KneeArrower* to retain the most significant interactions (1 698 edges). We visualised and analysed the resulting network with Cytoscape v3.10.2 (Shannon *et al*, 2003).

## Results

We used data from 300 patients with severe obesity who underwent bariatric surgery from the ABOS cohort. The current study relies on clinical and anthropometric data, RNA-seq from liver tissue, and CpG methylation from liver tissue (measured by Illumina EPIC arrays).

### Clinical data

Sixty-six clinical, medical, and anthropometric measurements were available for the patients, including blood labs, hypertension, diabetes, and dyslipidaemia status and treatment. We retained 16 non-redundant labs relevant for NAFLD’s clinical presentation, plus the sex and patient age for our clinical data module (Table 1). Missing values were imputed (See Materials and Methods).

A principal component analysis (PCA) showed that the disease status (i.e., {0, 1, 2} = {healthy, NAFL, NASH}) significantly aligned mostly with PC1 (Figure 1A), representing 19.6 % of the variance (R = −0.50, p = 3.45·10^−17^). Age, a known confounding factor for liver disease (McPherson *et al*, 2017), aligned partly with PC1 (R = −0.38, p = 6.17·10^−10^) but mainly with PC2 representing 11.8 % of the variance (R = 0.56 p = 1.38·10^−21^). Sex significantly aligned with PC1, but mostly with PC5 (7.8 % of the variance).

**Figure 1:**
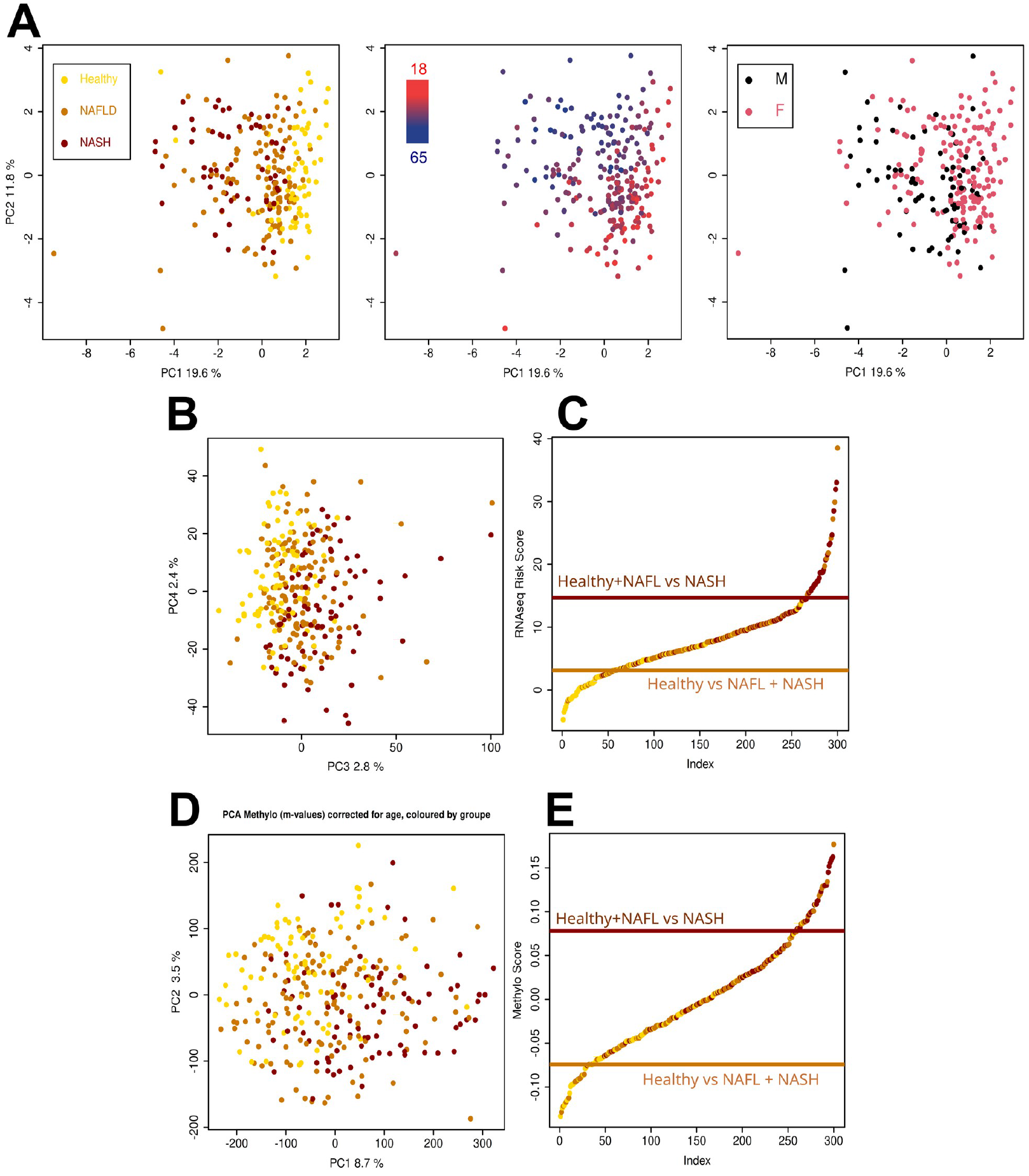
Features used to train the models. A) PCA of the retained 18 clinical variables. Severity (left) aligns mainly with PC1, while age aligns mostly with PC2. B) 3^rd^ and 4^th^ principal components of the RNA-seq PCA coloured by disease status. C) Ranked RNA scores for each subject determined with 2×100 main loadings, coloured by disease status. Horizontal lines represent the thresholds determined by logistic regression. D) First two components of the methylation m-values after correction of CpG methylation for age coloured by disease status. E) Ranked methylation scores for each subject determined with 2×130 main loadings coloured by disease status.

### Liver transcriptomics

Gene expression in the liver was obtained by total RNA-seq for all participants. PCA of the gene expression showed that liver disease status significantly aligned with PC3, representing only 2.8 % of the variance (R = 0.46 p = 5.56·10^−17^) (Figure 1B). Age significantly aligned with PC5, representing less than 2 % of the variance (R = −0.30 p = 9.3·10^−8^). To decide how many genes to use in the deep learning model, we created a classifying score by multiplying each gene expression by its PC3 loading. We then summed those scores over *N* genes with extreme positive loadings (on the right, associated with a more severe liver disease status) and *N* genes with extreme negative loadings (on the left, associated with a healthier status). Thus, patients with highly expressed genes on the left have a very negative score and are classified as “healthy”, while patients with highly expressed genes on the right have a very positive score and are classified as severely diseased (note that negative and positive can be swapped, as often with PCA, without changing the outcome). Via logistic regressions, we determined the optimal scores separating 1) all NAFLD *versus* healthy and 2) healthy plus NAFL *versus* NASH (shown for N = 100 on Figure 1C). To evaluate the number of genes to retain, we ran this procedure for N = 1 to N = 30 000 starting from the two extreme scores. After N = 100 transcripts, the accuracy decreased. The retained 200 transcripts (corresponding to 200 different genes) can be found in Supplementary Table 1, sheet “differential expression”. All these genes have a base mean expression over 2 TPMs.

### Liver DNA Methylation

We measured the DNA methylation of over 700 K CpGs in the liver with methylation arrays in all participants. A PCA of m-values showed that disease status aligned with PC1, representing 8.5 % of the variance (R = 0.48, p = 2.72·10^−18^) and with PC2, representing 6.5 % of the variance (R = −0.39, p = 2.17·10^−12^) (Supplementary Figure 1). CpG methylation is known to increase with age (Horvath, 2013), regardless of health status. Accordingly, it strongly aligned with PC2 (R = −0.72, p = 1.56·10^−49^). In order to select CpGs solely based on their relation to severity, we removed the age component. For each CpG, we fitted a linear model of methylation m-values in function of age and kept the residuals, producing a dataset where the age signal was completely removed. After this correction, the disease status still aligned with PC1 (Figure 1D), now representing 8,7 % of the variance (R = 0.39, p = 4.07·10^−18^), and with PC2, now representing only 3.5% % of the variance (R = −0.30, p = 7.91^−12^) confirming that NAFLD is associated with strong epigenetic remodelling in the liver. Since most of the variance was associated with PC1, we used the same procedure as for the RNA-seq to determine that 130 main loadings CpGs on each side of PC1 were optimal for the classification of disease status (Figure 1E, Supplementary Table 2, sheets “130PC1risk” and “130PC1protect”).

### Performance of the multi-module model

We built multi-layer perceptrons (MLPs) trained to recognize disease status from clinical data, gene expression, and DNA methylation, either individually or in combination (Figure 2). The best results were obtained with three modules reading the three data types. The difference with the models using only one or two of the datatypes is particularly marked when distinguishing NAFL from healthy liver and NASH (Figure 3A). We extensively explored different architectures, including the number of layers and of neurons per layer, activation functions, normalisation and regularisation, in conjunction with other hyper-parameters. The model architecture presented in Figure 2A provided the best results.

**Figure 2:**
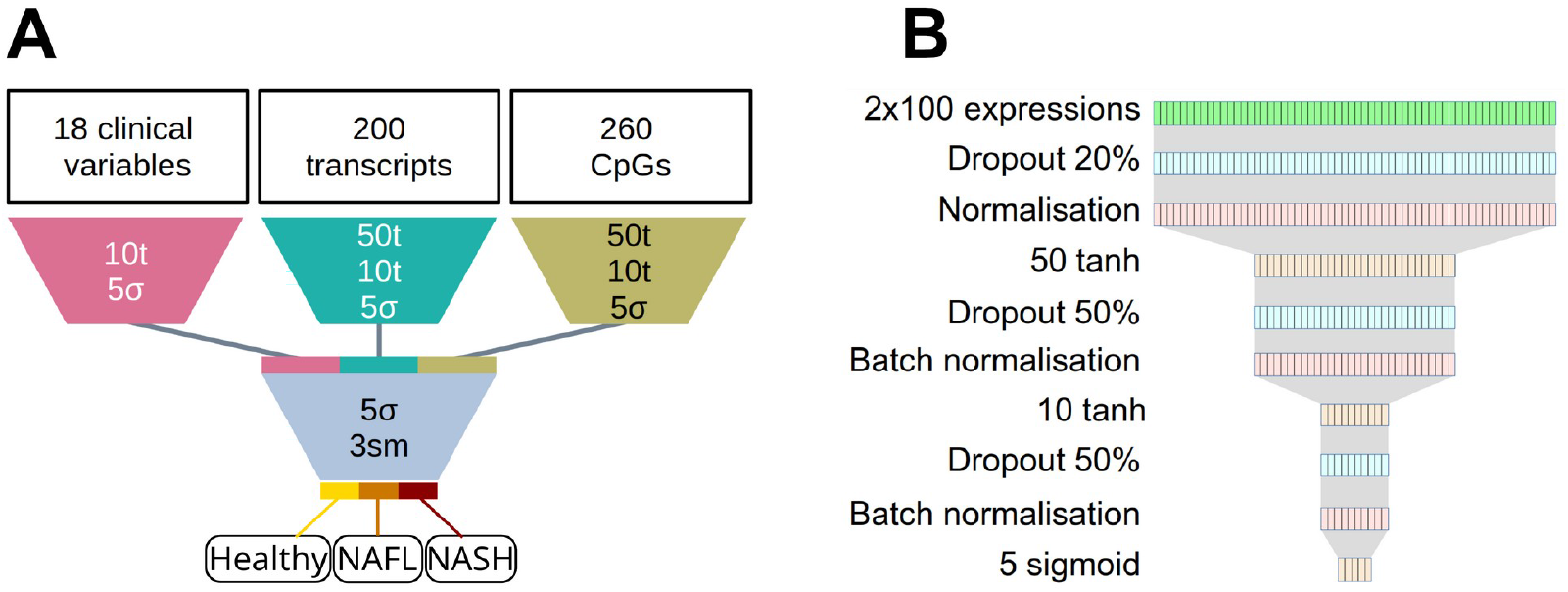
Architecture of the deep learning model. A) Global structure of the multi-module model. B) Detailed structure of the module reading RNA-seq data.

**Figure 3:**
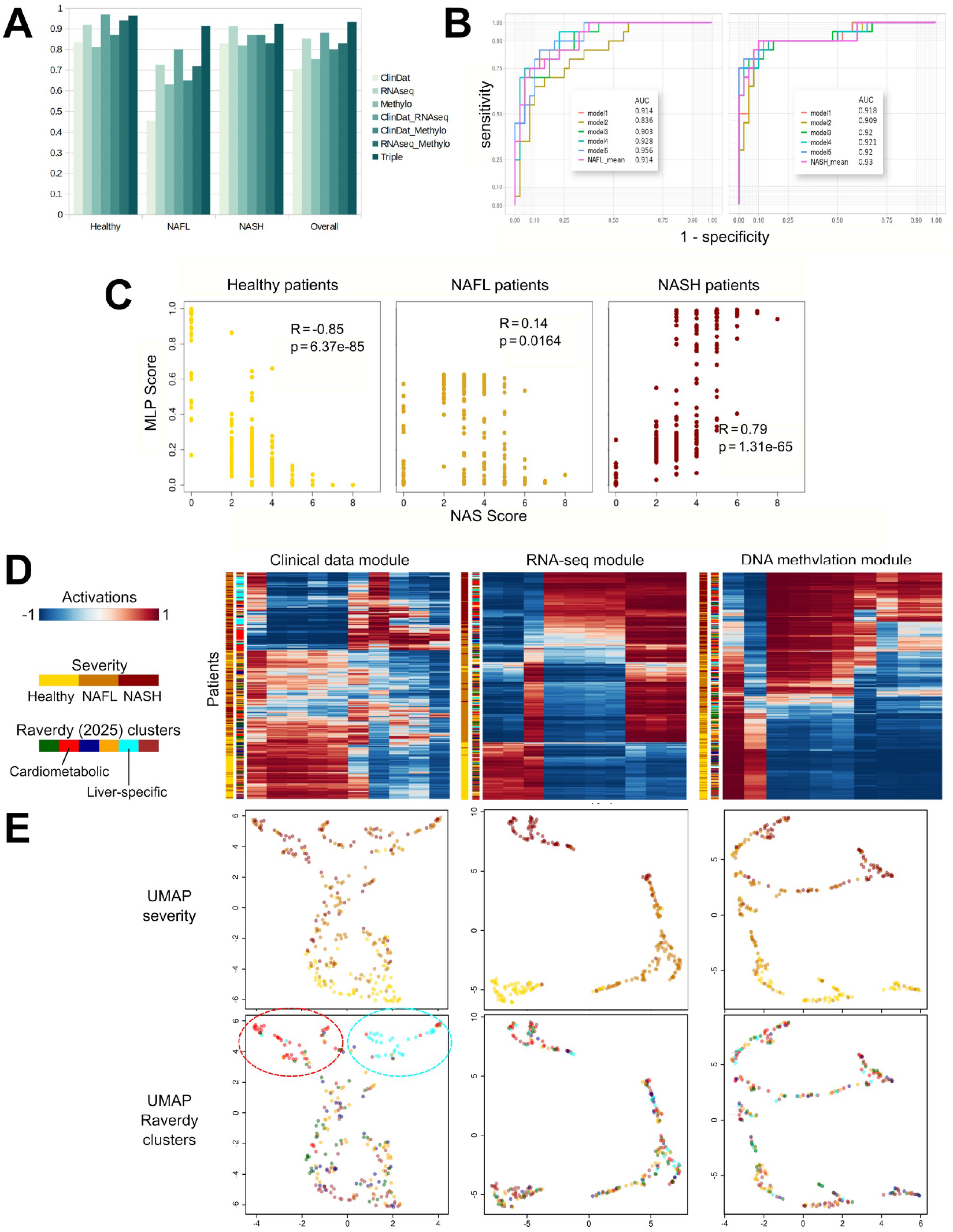
A) AUC of the different architectures, trained with one, two, or the three data types. B) ROC curve of the five independent models and the mean scoring for NAFL (left) and NASH (right) on the validation set. C) Correlation between the three prediction scores returned by the MLP and the NAS scores computed as the sum of the steatosis, ballooning, and inflammation scores. Left panel, healthy subjects; centre panel, NAFL patients; right panel, NASH patients. D) Clustering of patients in the latent space of each module (activations in the 10 tanh neuron layer of each module). Patients are coloured by severity and their belonging to the clusters defined by Raverdy et al 2025. Activations (columns) are also clustered. E) UMAP of neuron activations for each patient, coloured by severity class (top) and Raverdy *et al* 2025 clusters (bottom).

From the 298 retained subjects (see Materials and Methods), we randomly selected a validation set of 20 subjects from each of healthy, NASH and NAFL (60 subjects, 20%). The remaining set featured 79 healthy, 83 NASH, and 136 NAFL subjects. To balance the classes, we duplicated the first two to obtain 360 subjects. We trained five independent models with 90 subjects randomly selected for test and 270 used for learning. Hyper-parameters used in the learning phase are presented in the materials and methods section.

We averaged the softmax scores of all five models to produce an overall score for the three statuses. The highest score was retained as the predicted status. On the never-seen validation set, the combined models achieved an AUC of 0.914 for simple steatosis (three-class accuracy of 81.7%) and 0.93 for NASH (three-class accuracy of 83.3%) (Figure 3B). Keeping only the best three models increased slightly the AUC for simple steatosis, at 0.935 (The AUC for recognising healthy subjects also increases, from 0.963 to 0.975) due to better output scores, despite not changing the prediction results and the accuracy for either of the disease status or overall. As expected, the prediction score for healthy status correlates negatively with the NAFLD Activity Score (NAS) (Kleiner *et al*, 2005) computed as the sum of the histological steatosis, ballooning, and inflammation scores (R = −0.85, p-value = 6.40·10^−85^) while the prediction score for NASH status correlates positively with the NAS score (R = 0.79, p-value = 1.30·10^-65^) (Figure 3C).

### Impact of each module on the classification

To understand better the reasons behind the prediction accuracy, we looked at the clustering of subjects in the latent spaces of each module. Clustering the activations of the ante-penultimate layers before concatenation, containing 10 tanh neurons each, shows that the three modules are already able to differentiate between the degree of severity (Figure 3D, Supplementary Figure 2). However, the performance of the RNA-seq module is by far the best, followed by the DNA methylation module, and then the module reading clinical data, as observed from the AUCs (Figure 3A). From instance, the 10 tanh layer RNA-seq module of model 3 comprises 2 neurons specifically recognising healthy subjects, 1 recognising non-NASH subjects, 3 recognising both stages of NAFLD, and 4 recognising NASH. All five models behave similarly (Supplementary Figure 2). While able to predict the range of severity, the methylation modules do not comprise neurons neatly grouped by class prediction, but either two groups of neurones predicting low and high severity, or a continuum of severity predictions.

Patient embeddings by the clinical module permit to retrieve the different clusters proposed by Raverdy *et al* 2025, the “cardiometabolic” and “liver-specific” clusters being located in different subregions of the volume containing the patients with more severe disease status (Figure 3E). On the contrary, patients belonging to those clusters are not embedded in the same region of latent spaces built by the gene expression and DNA methylation modules, but are spread in the well-defined sub-regions containing healthy, NAFL, and NASH patients. Within the sub-regions, patients from both clusters did not segregate from each other. Differential gene expression between the cardiometabolic or the liver-specific cluster and control clusters (Supplementary Figure 3) showed a significant overlap with the genes differentially expressed between NAFLD and non-NAFLD patients (79% in both cases). However, this was much less the case for genes differentially expressed between the cardiometabolic and the liver-specific clusters 32/66 with NASH/Healthy and 18/66 with NAFL/Healthy. Only 7 genes differentially expressed between the cardiometabolic and liver-specific clusters were read by our gene expression module (*ATF3, JUN, MMP19, MMP9, NR4A1, NR4A3, SOCS3*).

To determine the relative contribution of each data type to the final classification, we analysed the connection weights between the concatenation layer, which integrates the outputs of the three modules, and the first merging layer. In all models, the synaptic weights with the largest absolute values were connecting the RNA-seq module to the merging layers, confirming that the chosen liver gene expressions contributed the most to the final classification (Figure 4A). The contribution of the other modules was nevertheless important since the RNA-seq module, on its own or in pair with either of the two other modules, provided less accurate classifications.

**Figure 4:**
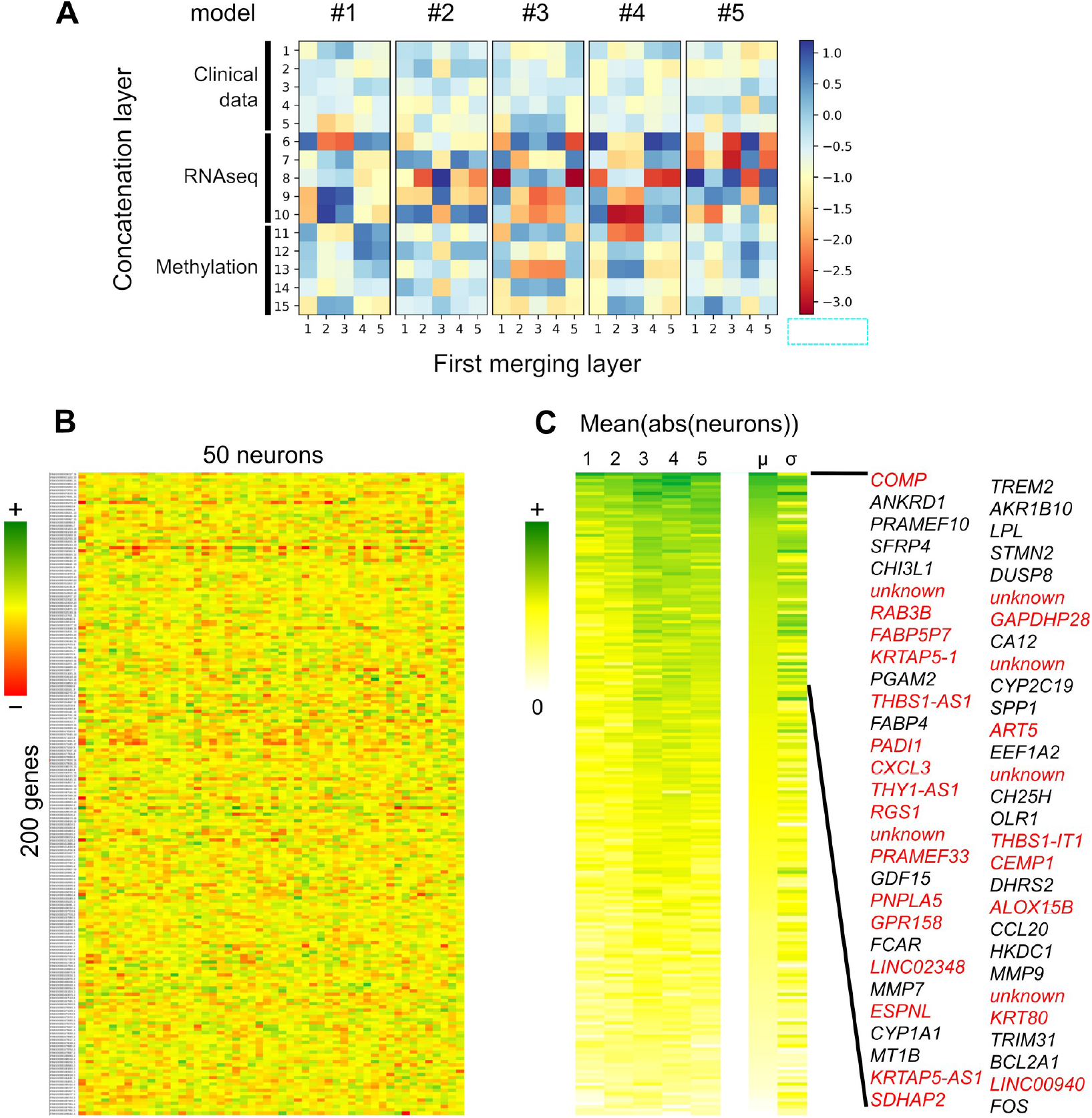
Synaptic weights of the RNA-seq module linking normalised gene expressions and the first layer of tanh neurons. A) Synaptic weighs between the concatenated output of the modules and the first merging layer. Colour intensity reflects the weights’ values and, thus the connection’s impact. B) all weights in model 1, ordered by Ensembl IDs; C) average of the absolute values of all 50 weights coming from each gene expression, for each of the five models, and their mean and standard deviation, ordered by absolute mean value. Black genes are known to be involved or markers of NAFLD while red ones are new.

### Exploration of the gene expression input

We compared the 200 genes selected for the RNA-seq module and the genes provided by differential expression analysis. Interestingly, some of the genes with the strongest loadings were not picked up by the differential expression analysis while known to be involved in NAFLD (e.g., *CXLC8* and *CCL20*). We found 212 genes differentially expressed between NAFL and healthy subjects, 558 between NASH and healthy subjects, and 63 between NASH and NAFL subjects. The 200 PC3 loadings are correlated with log2 fold changes between healthy and NASH subjects. However, many changes did not reach significance with differential expression analysis (Supplementary Figure 4, Supplementary Table 1, sheet “Differential expression”). Five of the 100 genes aligning the most with severity (left, positive loadings on PC3) were unannotated, and their function is unknown (Supplementary Table 1, sheet “PC3 loadings”). More strikingly, 39 of the 100 genes aligning the most with a healthy status were unannotated. Genes aligned with healthy status include insulin, reflecting the well-known links between liver disease and diabetes, and many long non-coding RNAs suggesting strong remodelling of regulatory networks.

To understand the relative impact of each gene expression on the final decision, we looked at the synaptic weights connecting the normalisation layer of the RNA-seq module and the first tanh neuron layer. Since tanh activations are symmetrical, a strong negative or positive input can have the same impact downstream through negative weights. Thus, we averaged the absolute values of the 50 weights connecting each gene expression to the downstream signal (Figure 4 right panel, Supplementary Table 3, sheet “Combined average(abs)”). The averaged values of different models showed a strong correlation, with R values ranging from 0.77 to 0.87 according to the pair of models (Supplementary Figure 5), showing that the five models learned to consider the same gene expressions. We then averaged the values across the five models to rank the gene expression impacts. Several of the most influential genes are involved in NAFLD or its complications (hepatic fibrosis, hepatocellular cancer), such as *TREM2* (2^nd^ highest absolute weight), *ANKRD1* (3^rd^), *AKR1B10* (4^th^), *LPL* (6^th^), *SFRP4* (7^th^) or markers such as *PRAMEF10* (5^th^), *STMN2* (8^th^), *CHI3L1* (9^th^). Many others have been found differentially expressed in the liver of NASH and healthy subjects. However, some of the most impactful genes have not been linked to NAFLD so far, such as *COMP* (which expression is multiplied by 25 in NASH compared to healthy) or *RAB3B* (which expression is multiplied by 2.4 in NASH compared to healthy).

We used a network inference approach to extend the list of 200 genes used in the gene expression module to the list of co-regulated genes. We used two independent network inference methods ( *CLR* and *GENIE3*) to compute a combined weight for interactions. After ranking them, we retained the interactions involving the 200 genes read by the gene expression module that have a rank above the “knee” (R package *KneeArrower*). This procedure resulted in a list 7 808 interactions involving 1 699 genes. The inferred interactions largely formed a mega-cluster, itself subdivided in a few highly connected sub-clusters (Figure 5), showing that the genes selected to train the models are largely co-regulated. The most highly connected genes (as per their degree) known to be involved in NAFLD included *DUSP8* (d = 166, 4^th^ rank), *CIDEC* (d = 122, 10^th^ rank), *UBD* (d = 118, 13^th^ rank), *STMN2* (d = 109, 19^th^), *TREM2* (d = 107, 22^nd^ rank), *GDF15* (d = 103, 25^th^ rank) (Supplementary Table 4). All these genes belonged to the main sub-cluster. The selection of all their first neighbours provided a list of 223 genes. A gene enrichment analysis with Gene Ontology Biological Process showed enrichments for terms linked with immune responses, largely due to chemokines (e.g. *Response to chemokine*, 12 genes, enrichment of 11-fold, FDR = 7.2·10^-7^), and extracellular matrix (e.g., *extracellular structure organization*, 17 genes, enrichment of 4.7-fold, FDR = 3.1·10^-5^) (Supplementary File, Supplementary Figure 6). Therefore, this largest sub-cluster points out that, unsurprisingly, inflammation and fibrosis are particularly important when the models decide which severity class a patient belongs to.

**Figure 5:**
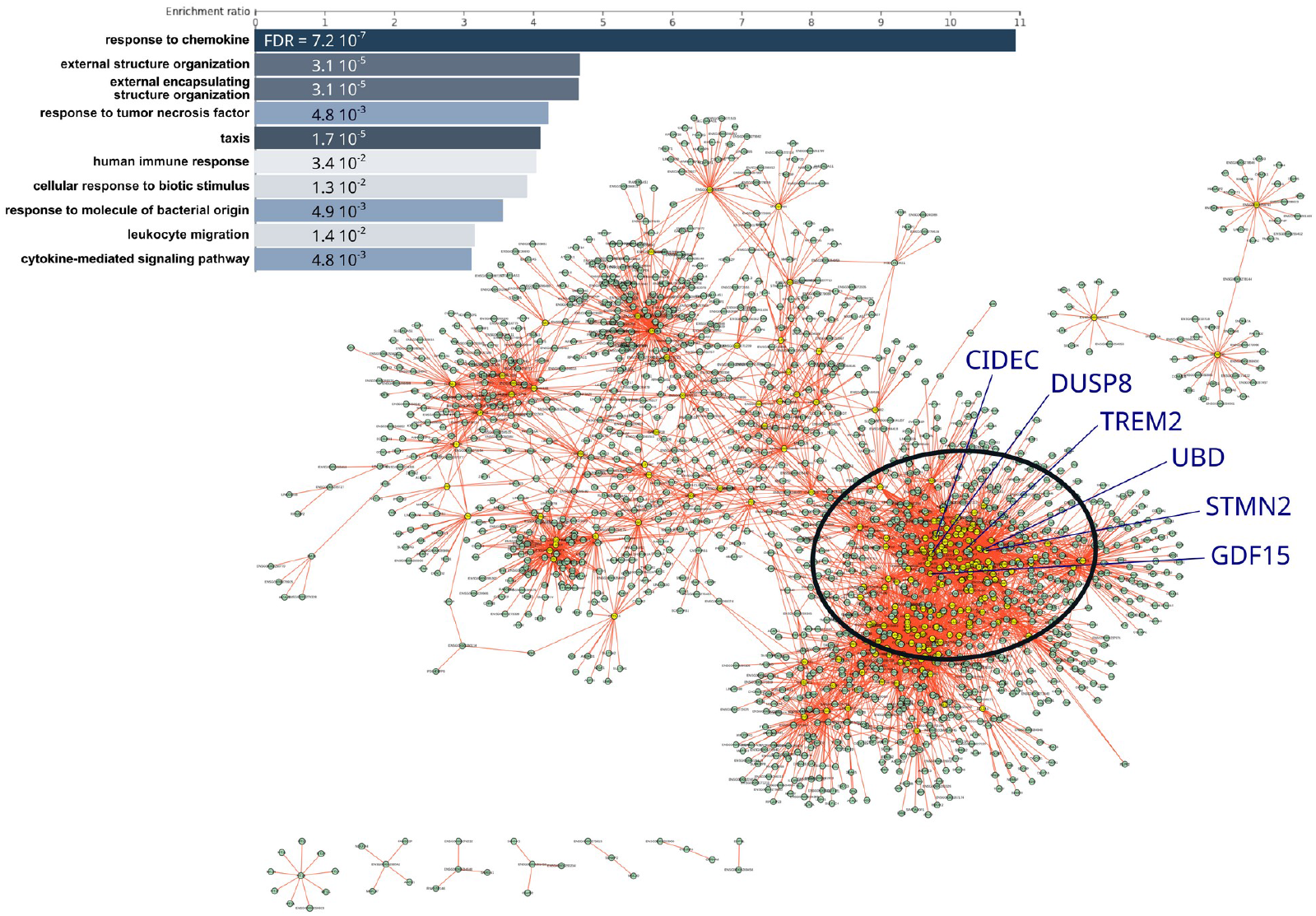
Interactions inferred from the RNA-seq dataset. We show the strongest interactions involving the 200 genes used to feed the RNA-seq MLP module. The highly connected nodes with a degree over 10 (i.e. connecting to over 10 other genes) are shown in yellow. The ellipse (bottom right) highlights the main subcluster, which members underwent a gene enrichment analysis (top left).

## Discussion

It is medically important to distinguish between isolated benign fatty liver and more severe MASH, and current approaches based on imaging or clinical data only are not effective enough, or are associated with serious side effects. To progress towards towards understanding the molecular underpinning of liver disease severity, we used both clinical and omics datasets to train deep learning models that recognise the disease status with a multi-class AUC of 0.945. In doing so, we identified genes whose expressions and methylation status are known to be involved in MASLD aetiology as well as new ones, worth investigating.

We extensively explored all hyperparameters and settled on tanh activation functions in each module, followed by sigmoid and softmax activation functions. Other combinations led to decreased accuracy. Decreasing the number of layers also decreased accuracy, while increasing the number and size of the layers led to overfitting. An important factor in increasing the accuracy was the inclusion of dropout (Hinton *et al*, 2012) and batch normalisation (Ioffe & Szegedy, 2015) layers after each non-linear layer. We created independent models learning to recognise the disease status from selected clinical data, gene expression, and CpG methylations, pairs of those datasets or the three together. The combination of five models comprising the three modules performed better overall, followed by the models reading clinical data plus RNA-seq, then those reading solely RNA-seq. The impact of using several modules is particularly noticeable on the prediction of “benign” steatosis. Models using clinical data alone did not recognise well liver steatosis (mean predictions with AUC of 0.455) despite recognising well liver healthy from NASH patients (mean predictions with AUC of 0.834 and 0.829 respectively).

Recently, Raverdy *et al* showed that using 6 clinical variables, patients from the studied cohort could be grouped in 6 different clusters (Raverdy *et al*, 2024). Two clusters showed a particular enrichment in patients with MASH. One cluster, dubbed “cardiometabolic” comprised patients who would later present a higher prevalence of cardiovascular diseases and type 2 diabetes, while a “liver-specific” cluster comprised patients who would later present lower prevalence of both diseases. This study’s module reading the clinical data is able to distinguish subjects belonging to the both clusters, embedded in sub-regions of the volume corresponding to more severe disease status (deeper principal components of a PCA permit to retrieve some of the other clusters too). However, patients belonging to the “cardiometabolic” and “liver-specific” clusters are not separated in the latent spaces of the RNAs-seq and DNA methylation modules. This is not surprising considering that only 7 of the 66 genes we found differentially expressed between the two clusters are part of the 200 genes whose expression is read by our model. As shown in Raverdy et al 2025, both clusters are enriched in NASH patients. However, the proportion of steatohepatitis is not what characterises them but rather the long-term evolution of non-hepatic diseases (Raverdy *et al*, 2024). The corresponding molecular endotypes are therefore likely different from the endotypes characterising the severity classes.

Many of the genes whose expressions correlate the most with disease severity (29 of the 35 highest PC3 loadings) – and thus selected for the RNA-seq module, are known to be involved in MASLD or related complications (including diabetes) – whether transcription factors (e.g., *FOSB, EGR3, SFRP4, ATF3, FOS, EGR2, NR4A1, OSM, NR4A3*), cytokines (e.g., *CXCL9* [IL-8], *CCL20, CXCL1, CCL2*), enzymes linked to lipid metabolism (e.g., *AKR1B10, LPL*), or signalling proteins (e.g., *DUSP8, SOCS3, AREG, ORL1* [LOX-1], *PTGS2* [COX-2], *CIDEC*) but also other types of proteins such as insulin, *UBD* (FAT10) and *GDF15*. Many of these genes, (e.g., *UBD* and *SPP1)* were identified as signatures of severity in men and women, and in animal models (Vandel *et al*, 2021). However, the functions of 5 of the 100 genes aligning the most with severity and 39 of the 100 genes aligning the most with a healthy status are unknown, pointing to novel avenues to explore the molecular networks underpinning liver pathophysiology and possibly discover new molecular therapy avenues. The average of absolute values of the weights connecting the RNA-seq input to the first non-linear layer also provides a ranked list of impacts. Interestingly, while most of the genes mentioned above are on the top part of this list, the correlation coefficient between PC3 loadings and the mean of absolute weights for the top 100 genes is only 0.48 (and even 0.27 for the entire set of 200) (Supplementary Table 3, sheet “Correlation weights-loadings”). This might indicate a difference between the amplitude of the expression difference and the actual impact on the severity. At the very least, it provides a new list of genes to investigate.

Zhang et al. proposed to use the liver expression level of four genes to distinguish NASH patients from controls (Zhang *et al*, 2023). Only one was common with our 200, *FOSB*, our strongest hit. From liver transcriptomics, Park et al derived a list of 10 genes to distinguish NASH from NAFL with a small 2-layer network of 24 neurons reaching an AUC of 0.805 on a validation set (Park *et al*, 2023). Their input list presented only three genes in common with ours (*AKR1B10, DTNA, MAST1*), although *BCAT1, FABP4, FABP5*, and *RGCC* – also in our list – were used to predict evolution towards hepatocellular cancer. Oh et al. derived a list of six signature genes, two of them – *TREM2* and *SPP1* – being also read by our RNA-seq module. Their liver expression allowed to discriminate, with an AUC of 0.86, steatosis from MASH with bridging fibrosis or cirrhosis, and, with an AUC of 0.767, steatosis from MASH without or with simple fibrosis (Oh *et al*, 2024). Larger panels of signature genes were also proposed to correlate with the severity of MASLD. Hoang et al. proposed 20 genes which form clusters whose liver expression profiles are characteristic of NAS levels (Hoang *et al*, 2019). *B9D1, CCL20, FABP5, PRAMEF10, TREM2*, and *UBD* are also on our list. Similarly, Govaere et al proposed 24 genes, of which *AKR1B10, CCL20*,33 *DTNA, DUSP8, GDF15, STMN2, THY1, TNFRSF12A* are also in our list, that provide AUC of 0.85 for discriminating NAS > 4 (Govaere *et al*, 2020). With AUC over 0.91 for both NAFL and NASH, our classifier outperformed the previous best models by around 10%, reducing errors by a third.

Collecting liver tissue for RNA-seq is invasive, and cannot be used to detect or grade MASLD routinely. The quest for non-invasive biomarkers of the disease severity is thus active. However, most non-omics blood biomarkers or panels of biomarkers currently yield AUCs of 0.7 to 0.85, review in (Piazzolla & Mangia, 2020; Zoncapè *et al*, 2024; Sanyal *et al*, 2023). Nevertheless, pioneering studies have started to use DNA methylation of whole blood or saliva to diagnose fibrosis (Sun *et al*, 2022) or hepatocellular carcinoma (Mezzacappa *et al*, 2024). Such approaches might allow the detection of differential liver DNA methylation in MASLD via circulating DNA..

In conclusion, we showed that deep learning models comprising modules reading clinical data, gene expression, and CpG methylations are able to recognise liver steatosis without inflammation from steatohepatitis with excellent accuracy. Analysing the respective impact of the various genes and CpG included in the models suggests new possible genes and molecular circuits involved in MASLD progression and severity. Although using molecular profiles from liver is not suitable for MASLD diagnosis, future deep-learning models reading selected clinical and multi-omics data from blood or saliva might provide powerful element for the diagnosis and prognosis toolkits.

## Supporting information

RNA-seq input

DNA methylation input

Weights 1st layer RNA-seq

Gene interactions

Gene enrichment

## Abbreviations

ABOS: Atlas Biologique de l’Obésité Sévère
AUC: area under the (ROC) curve
CLR: context likelihood of relatedness
TPM: transcript per million readss
CRP: c-reactive protein
GENIE3: gene network inference with ensemble of trees
HDL: high-density lipoproteins
kNN: k nearest neighbours
LDL: low-density lipoproteins
MAFL: metabolic dysfunction–associated steatotic liver
MASH: metabolic dysfunction associated steatohepatitis
MASLD: metabolic dysfunction associated steatotic liver disease
MLP: multi-layer perceptron
NAS: NAFLD activity score
NASH: non-alcoholic steatohepatitis
NAFL: non-alcoholic fatty liver
NAFLD: non-alcoholic fatty liver disease
OGTT: Oral Glucose Tolerance Test
PCA: principal component analysis
ROC: receiver operating characteristic

## Data availability statement

The code to build, train and use the models presented in this study is available at https://gitlab.com/ngamblen/mlp-precinash

## Acknowledgments

Authors thank Anne-Sophie Ledoux, Vincent Massy et Frédéric Allegaert for their technical support. This study was funded by the National Center for Precision Diabetic Medicine – PreciDIAB (ANR-18-IBHU-0001; FEDER 20001891/NP0025517; MEL 2019_ESR_11), the Agence Nationale de la Recherche (ANR) grants PreciNASH (16-RHUS-0006) and European Genomic Institute for Diabetes (Egid, 10-LABX-0046). The study was also supported by a Contrat de Plan État-Régions (CPER) from the Conseil Régional des Hauts-de-France (N° 2020-R3-CTRL_IPL_Phase4), and the program Investissements d′Avenir I-SITE ULNE / ANR-16-IDEX-0004 ULNE as well as Horizon Europe Research and Innovation Programme OBELISK (101080465). NG and MA benefits from a STaRS funding from rom the Conseil Régional des Hauts-de-France. SF is funded by PreciDIAB. AB is supported by the European Research Council (OpiO 101043671).

