## Supplementary figures for "Deep learning models reading clinical data and liver omics strongly distinguish NASH from steatosis and suggest new genes involved in liver disease severity"

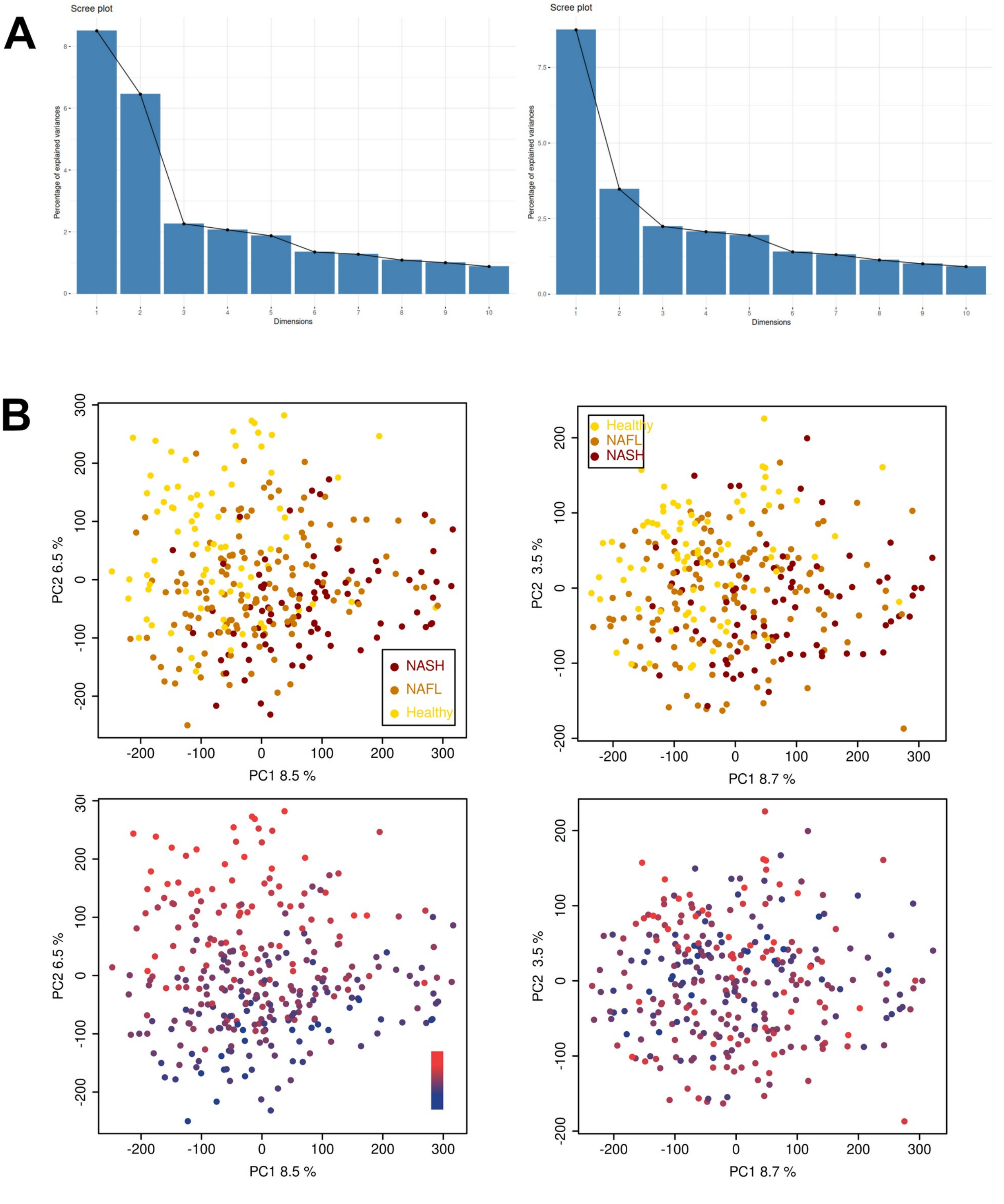


**Supplementary figure 1:** A)Variances associated with the first 10 principal components of a PCA ran on CpG methylation m-values, before (left) and after (right) correction for age. B) First two principal components of the CpG methylation m-values PCA before (left) and after (right) correction for age, coloured by severity (top) or age (bottom).

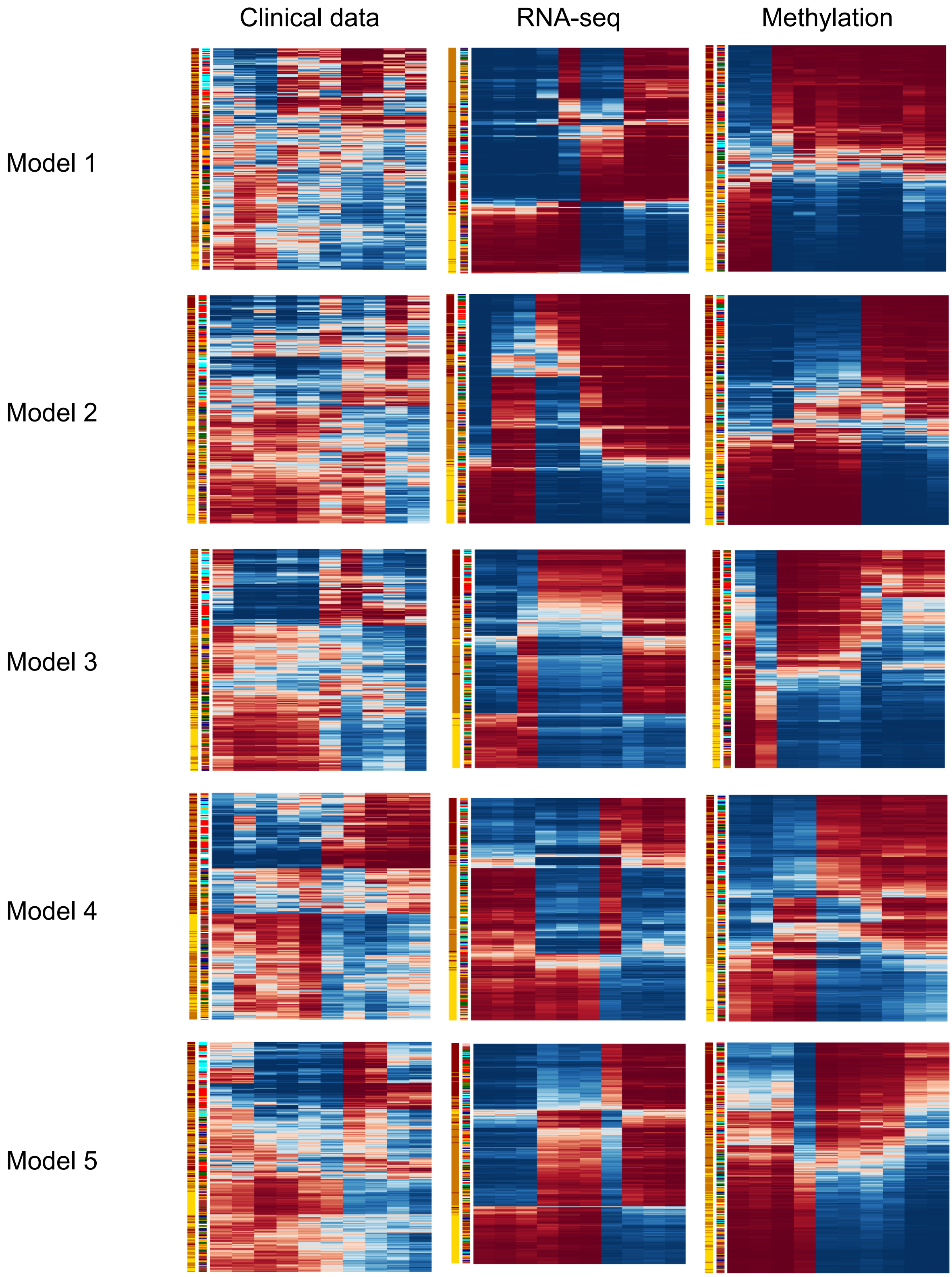


**Supplementary Figure 2**: Clustering of activations in the 10 tanh neuron layer of each module. Patients are coloured by severity and their belonging to the clusters defined by Raverdy et al 2025. Columns are clustered neurons neurons and rows are clustered patients.

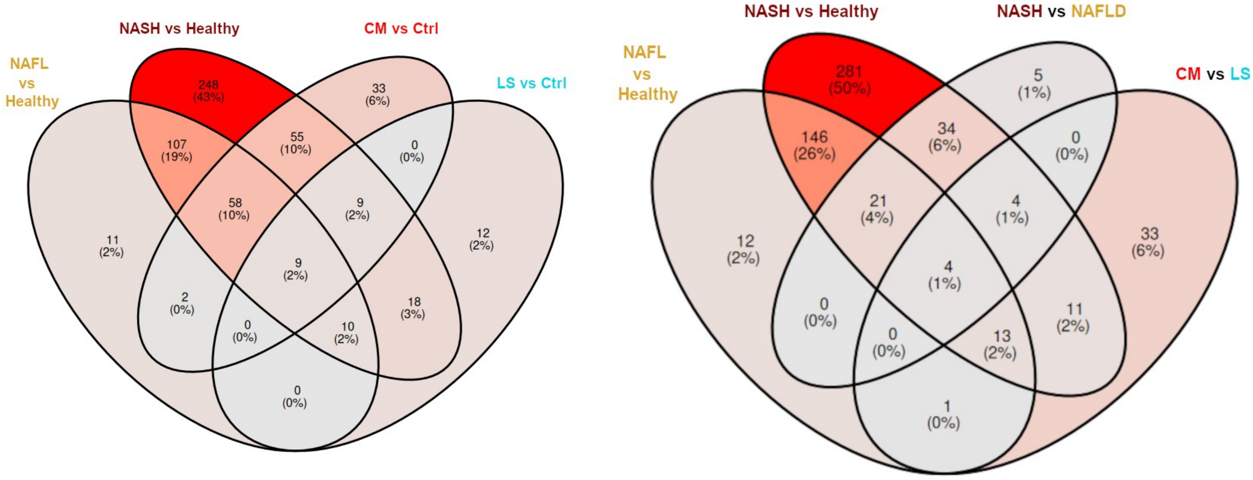


**Supplementary Figure 3**: Venn diagrams showing the number of genes differentially expressed between severity status or between clusters defined in Raverdy *et al* 2024. As in the latter paper, the group "Ctrl" is formed by merging their clusters 1, 3, and 6.

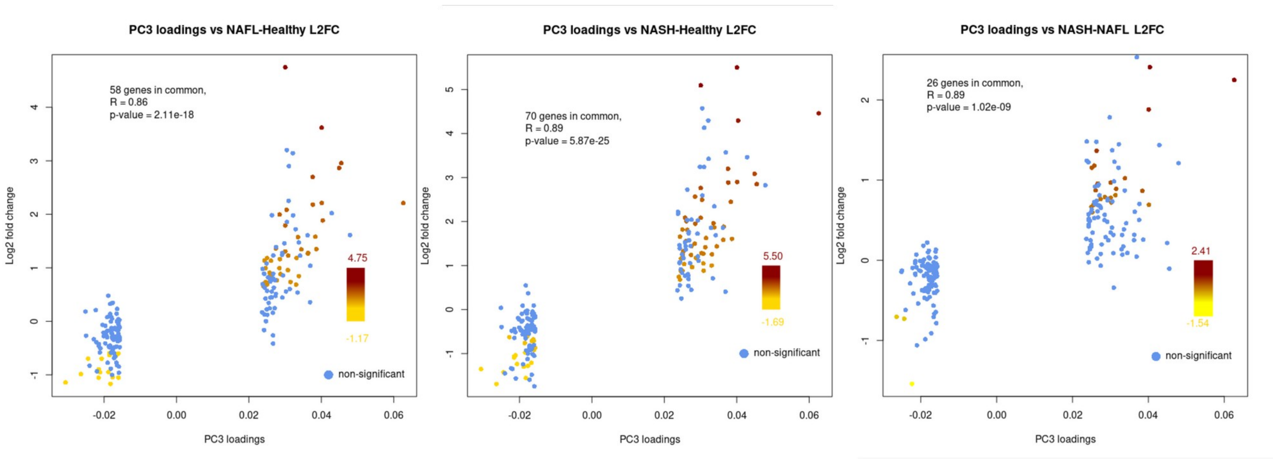


**Supplementary Figure 4**: Correlation between the PC3 loadings and the log fold changes between severity statuses. Blue dots represent genes present in the subset selected from RNA-seq PC3 that are not found significant by DESeq2. The other dots represent genes present in the subset selected from RNA-seq PC3 that are found significant by DESeq2, coloured by log3 fold change. The blank discontinuity in the middle of loadings comes from the selection of only the 100 more extreme values on each side of PC3.

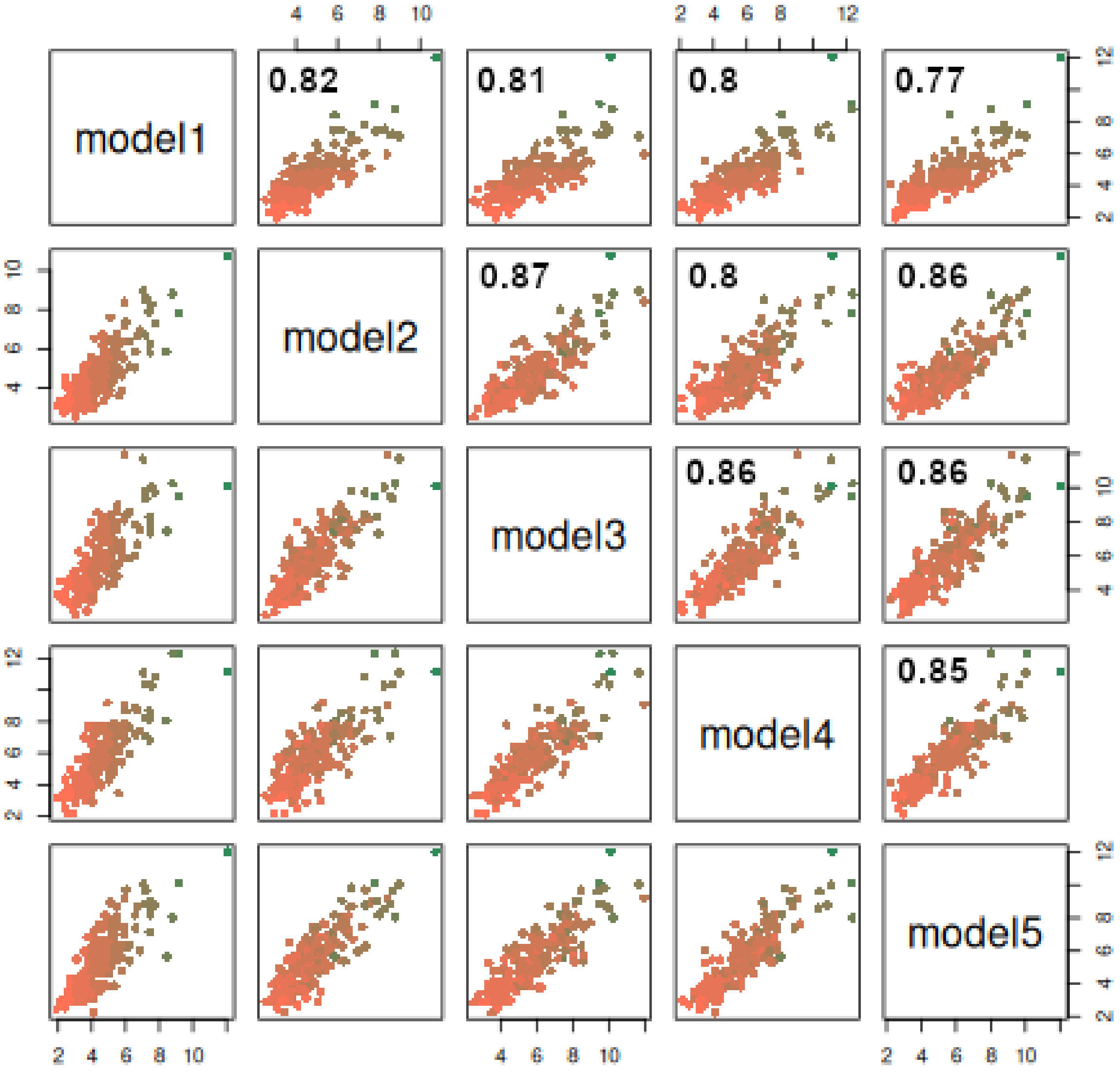


**Supplementary Figure 5**: Comparison of the absolute values of the weights connecting the normalised RNA-seq inputs and the first tanh layer. Pearson correlation coefficients between pairs of model are indicated on the top panels.

Select an enriched analyte set...

GO:1990868: response to chemokine




**Analyte set: GO:1990868 response to chemokine**

| FDR | P Value | | |
|---|---|---|---|
| 7.2147e-7 | 8.5992e-10 | | |
| **Analyte Set Size** | **Expected Value** | **Overlap** | **Enrichment Ratio** |
| 90 | 1.0969 | 12 | 10.940 |

| User ID | Gene Symbol | Gene Name | Entrez Gene ID |
|---|---|---|---|
| ACKR3 | ACKR3 | atypical chemokine receptor 3 | 57007 |
| CCL2 | CCL2 | C-C motif chemokine ligand 2 | 6347 |
| CCL20 | CCL20 | C-C motif chemokine ligand 20 | 6364 |
| CCL22 | CCL22 | C-C motif chemokine ligand 22 | 6367 |
| CXCL1 | CXCL1 | C-X-C motif chemokine ligand 1 | 2919 |
| CXCL10 | CXCL10 | C-X-C motif chemokine ligand 10 | 3627 |
| CXCL6 | CXCL6 | C-X-C motif chemokine ligand 6 | 6372 |
| CXCL8 | CXCL8 | C-X-C motif chemokine ligand 8 | 3576 |
| CXCL9 | CXCL9 | C-X-C motif chemokine ligand 9 | 4283 |
| PF4V1 | PF4V1 | platelet factor 4 variant 1 | 5197 |
| TREM2 | TREM2 | triggering receptor expressed on myeloid cells 2 | 54209 |
| ZC3H12A | ZC3H12A | zinc finger CCCH-type containing 12A | 80149 |

20 per page

1

Select an enriched analyte set...

GO:0043062: extracellular structure organizatio

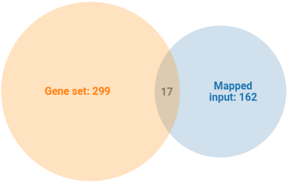


**Analyte set: GO:0043062 extracellular structure organization**

| FDR | P Value | | |
|---|---|---|---|
| 0.000030745 | 1.3973e-7 | | |
| **Analyte Set Size** | **Expected Value** | **Overlap** | **Enrichment Ratio** |
| 299 | 3.6441 | 17 | 4.6650 |

| User ID | Gene Symbol | Gene Name | Entrez Gene ID |
|---|---|---|---|
| ADAMTS4 | ADAMTS4 | ADAM metallopeptidase with thrombospondin type 1 motif 4 | 9507 |
| AEBP1 | AEBP1 | AE binding protein 1 | 165 |
| ANTXR1 | ANTXR1 | ANTXR cell adhesion molecule 1 | 84168 |
| ANXA2 | ANXA2 | annexin A2 | 302 |
| CAPG | CAPG | capping actin protein, gelsolin like | 822 |
| CCDC80 | CCDC80 | coiled-coil domain containing 80 | 151887 |
| COL14A1 | COL14A1 | collagen type XIV alpha 1 chain | 7373 |
| COL1A1 | COL1A1 | collagen type I alpha 1 chain | 1277 |
| COL3A1 | COL3A1 | collagen type III alpha 1 chain | 1281 |
| COMP | COMP | cartilage oligomeric matrix protein | 1311 |
| ELF3 | ELF3 | E74 like ETS transcription factor 3 | 1999 |
| LAMA2 | LAMA2 | laminin subunit alpha 2 | 3908 |
| LOXL1 | LOXL1 | lysyl oxidase like 1 | 4016 |
| LOXL4 | LOXL4 | lysyl oxidase like 4 | 84171 |
| LUM | LUM | lumican | 4060 |
| MMP7 | MMP7 | matrix metallopeptidase 7 | 4316 |
| MMP9 | MMP9 | matrix metallopeptidase 9 | 4318 |

**Supplementary figure 6**: Two most enriched Gene Ontology Biological Process terms in the 223 members of the neighbourhood of {*DUSP8, CIDEC, UBD, STMN2, TREM2, GDF15*}, with the relevant annotated genes.
