## Supplementary figures and images for "Deep learning models reading clinical data and liver omics strongly distinguish NASH from steatosis and suggest new genes involved in liver disease severity"

### goslim_summary_wg_result1722931927.png

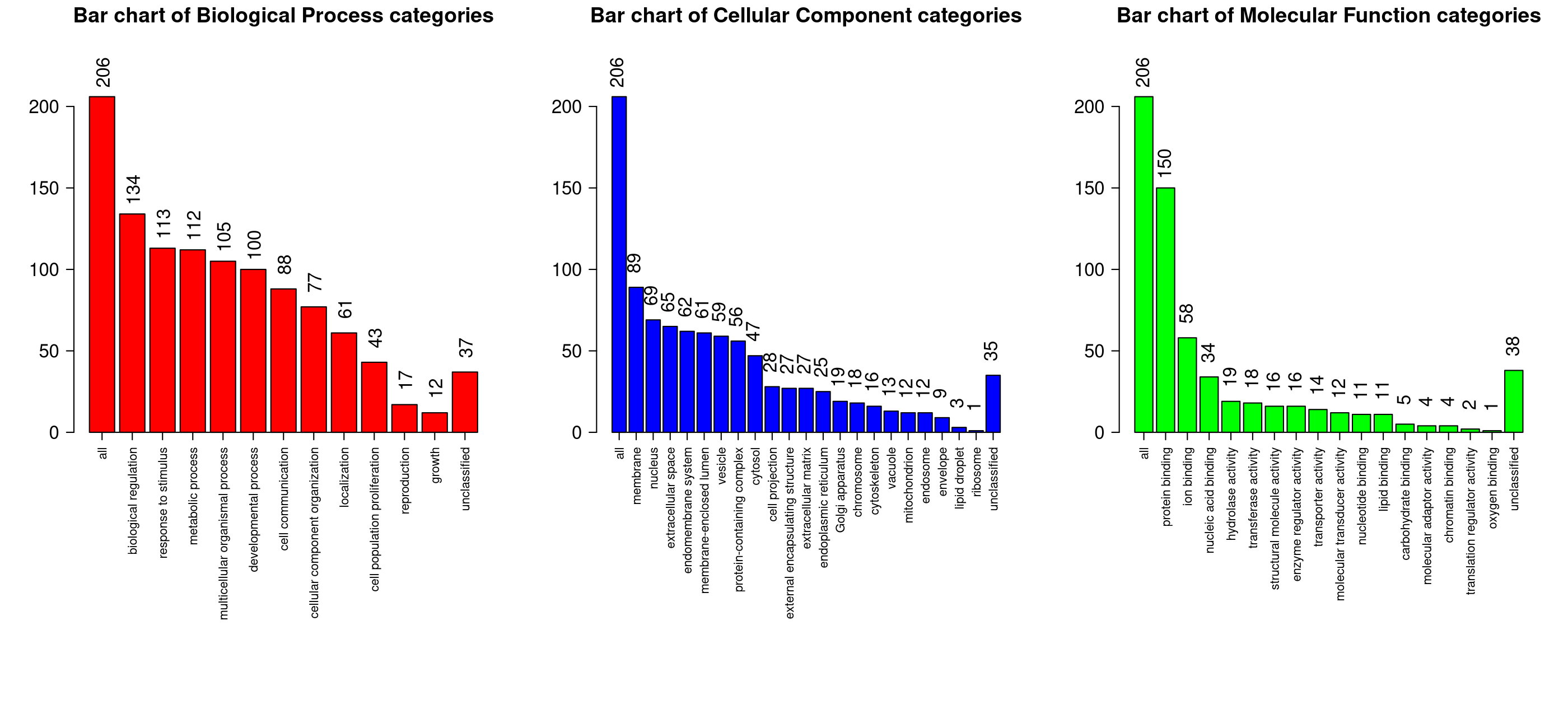
